# Tracing foot-and-mouth disease virus phylogeographical patterns and transmission dynamics

**DOI:** 10.1101/590612

**Authors:** Manuel Jara, Alba Frias-De-Diego, Simon Dellicour, Guy Baele, Gustavo Machado

## Abstract

Foot-and-mouth disease virus (FMDV) has proven its potential to propagate across local and international borders on numerous occasions, but yet details about the directionality of the spread along with the role of the different host in transmission remain unexplored. To elucidate FMDV global spread characteristics, we studied the spatiotemporal phylodynamics of serotypes O, A, Asia1, SAT1, SAT2, and SAT3, based on more than 50 years of phylogenetic and epidemiological information. Our results revealed phylogeographic patterns, dispersal rates, and the role of host species in the dispersal and maintenance of virus circulation. Contrary to previous studies, our results showed that three serotypes were monophyletic (O, A, and Asia1), while all SATs serotypes did not evidence a defined common ancestor. Root state posterior probability (RSPP) analysis suggested Belgium as the country of origin for serotype O (RSPP=0.27). India was the ancestral country for serotypes A (RSPP= 0.28), and Asia-1 (RSPP= 0.34), while Uganda appeared as the most likely origin country of all SAT serotypes (RSPP> 0.45). Furthermore, we identified the key centers of dispersal of the virus, being China, India and Uganda the most important ones. Bayes factor analysis revealed cattle as the major source of the virus for most of the serotypes (RSPP> 0.63), where the most important host-species transition route for serotypes O, A, and Asia1 was from cattle *Bos taurus* to swine *Sus scrofa domesticus* (BF>500), while, for SAT serotypes was from *B. taurus* to African buffalo *Syncerus caffer*. This study provides significant insights into the spatiotemporal dynamics of the global circulation of FMDV serotypes, by characterizing the viral routes of spread at serotype level, especially uncovering the importance of host species for each serotype in the evolution and spread of FMDV which further improve future decisions for more efficient control and eradication.

## INTRODUCTION

The rapid growth of global population along with the current demand for animal protein and the increasing animal trade have increased the spread of a broad range of transboundary animal diseases (TADs) [1, 2]. Clear examples of this phenomenon are the recent emergence of African Swine Fever in Asia, Avian Influenza or Contagious Bovine Pleuropneumonia and the return of Foot-and-mouth disease virus (FMDV) to Korea and other countries which successfully eliminated the virus for several years [3–7]. Understanding the tempo and mode of disease evolution allows to estimate the impact of external factors influencing these diseases and assess the evolutionary patterns followed over time, which can be better studied by considering the most recent advances in virus sequencing and phylogenetics [8–10].

Molecular phylogenetics has shown to be an accurate and highly impacting approach in the understanding of the spatiotemporal dynamics of infectious diseases, with the capacity to explain disease spread, virulence and invasion potential [11–20]. In the case of TADs, molecular phylogenetics has also proven to be a useful approach, providing accurate knowledge for the control of pathogens worldwide [21–24], however, it remains underused.

FMDV causes the most influential transboundary animal disease with historical worldwide circulation reported in domestic and wildlife reservoirs [25, 26]. FMD is a highly contagious disease caused by a small single-stranded RNA virus of the genus *Aphthovirus*, member of the family Picornaviridae [27] and classified into seven different serotypes; O, A, C, Asia1 and Southern African Territories (SATs) 1, 2 and 3 [25, 28–30] which severely affect the productivity of domesticated livestock, causing great economic losses [31–33]. The United States Department of Agriculture has estimated that the introduction of FMDV could result in losses between $15 to $100 billion [34–37]. One of the main reasons for this great impact is the wide variety of hosts known for FMDV (i.e., cattle, buffalo, swine, sheep, and deer), altogether it affects more than 70 species of cloven-hoofed animals [38]. FMDV is known to be transmitted locally and globally, often associated with infected animal products and human and animal movements [39, 40]. Transmission is also facilitated by airborne spread, direct animal contact with infected individuals or carcasses and translocation of contaminated staff, equipment, and machinery [41, 42].

Individual genes have been widely used to study the phylogenetic relationships among FMD serotypes [43–50], however, little has been done using whole genome sequences (WGS) [26, 51–53]. Nevertheless, these studies have been often based on a limited number of sequences (<200), or on phylogenetic methods that do not have the ability to accommodate uncertainty (i.e., Bayesian phylodynamic methods) [54]. Although these methods have been widely used, they present certain degree of evolutionary inaccuracy since in most cases (excluding Yoon et al (2011) and Omondi et al (2019)) the Bayesian phylogenetic and phylodynamic methods were not considered. Thus, their inherent ability to accommodate uncertainty, and therefore to assess the level of error of the predictions obtained, was often neglected.

Several studies have explored the spatiotemporal evolutionary dynamics of FMDV in different parts of the world, mainly focusing on the diffusion patterns across its endemic regions: Asia and Africa [31, 48, 55–59] However, studies considering all serotypes are only available within small geographic regions, and global studies only assessed some of the viral serotypes [56, 60].

In this study, we investigated the spatiotemporal dynamics of FMDV by using Bayesian phylodynamic analyses of comprehensive genetic, geographical and temporal data regarding past FMDV occurrences. The objectives of this work were to reconstruct the global evolutionary epidemiology of FMDV serotypes O, A, Asia1, and SATs, to make comparisons among the global spatiotemporal spread of each FMDV serotype, identify ancestral countries and provide inferences about the evolutionary patterns and the transmission between host species.

## MATERIALS AND METHODS

### Data collection and curation

We built a comprehensive genetic database comprising 249 publicly available whole genome sequences from six FMDV serotypes (A, O, Asia1, SAT 1, SAT 2 and SAT 3), with collection dates ranging from 1959 to 2017 (GenBank ID numbers in Supplementary material Table S1). Serotype C was not included in this study due to data unavailability (only 3 sequences were available). Our dataset gathers information from 43 countries and 4 continents obtained from the Virus Pathogen Resource database, available at https://www.viprbrc.org (See Table S1). To determine accurate phylogenetic relationships among FMDV reports, we combined the available genetic information along with collection date, host species (i.e., *Bos taurus*= cattle, *Syncerus caffer*= African buffalo, *Bubalus bubalis*= Water buffalo, *Sus scrofa domesticus*= swine, *Sus scrofa*= boar, and *Ovis aries*= sheep) and location (discrete information at country level) as metadata information. Any sample lacking one of these three characteristics was discarded.

### Discrete phylogeographical analysis

Sequences were aligned using Mega X, available at www.megasoftware.net [61]. The recombination detection program (RDP) v5.3 was used to search for evidence of recombination within our dataset [62]. Each serotype was screened using five different methods (BootScan, Chimaera, MaxChi, RDP, and SiScan). After removing all the duplicated sequences (i.e., representing the same outbreak), no evidence of recombinant sequences was observed in any FMDV serotypes analyzed. To determine whether there was a sufficient temporal molecular evolutionary signal of the FMDV sequences used for each serotype phylogeny, we used TempEst v1.5 [63]. To calculate the *P*-values associated with the phylogenetic signal analysis, we used the approach described by [64] based on 1,000 random permutations of the sequence sampling dates [65]. The relationship found between root-to-tip divergence and sampling dates (years) supported the use of molecular clock analysis in this study for all serotypes. Root-to-tip regression results for each serotype are reported in Supplementary Table S2, all the results supported a significant temporal signal (*P*-value<0.05). Phylogeographic history of FMDV dispersal was recovered from the obtained spatiotemporal phylogenies for each serotype. Phylogenetic trees were generated by a discrete phylogeography estimation by Bayesian inference through Markov chain Monte Carlo (MCMC), implemented in BEAST v2.5.0 [66]. We partitioned the coding genes into first+second and third codon positions and applied a separate Hasegawa-Kishino-Yano (HKY+G; [67]) substitution model with gamma-distributed rate heterogeneity among sites to each partition [68].

By using Nested Sampling Beast package v1.0.4 [69] we compared different molecular clock models to find the one that showed the best fit for the data related to each serotype. The marginal likelihood value supported the use of uncorrelated lognormal relaxed molecular clock [70]. To infer the epidemic demographic histories of FMDV per each serotype we estimated the effective number of infections through time by using the Bayesian skyline plot approach [71]. All analyses were developed for 200 million generations, sampling every 10,000^th^ generation and removing 10% as chain burn-in. All the Markov chain Monte Carlo analyses for each serotype were investigated using Tracer software v1.7 [72] to ensure adequate effective sample sizes (ESS) (above 200), which were obtained for all parameters. Final trees were summarized and visualized via Tree Annotator v. 2.3.0 and FigTree 1.4.3 respectively (included in BEAST v2.5.0) [66, 73].

To reconstruct the ancestral-state phylogeographic transmission across countries and hosts, we used the discrete-trait extension implemented in BEASTv2.5.0 [66]. In addition, to explore the most important historical dispersal routes for the spread FMDV across countries, as well as most probable host-species transition, we used a Bayesian stochastic search variable selection (BSSVS) [74]. Using BSSVS approach, we identified and eliminated the nonzero rates of change between each pair of discrete traits (countries and hosts species) based on its Bayes factor value obtained (lower than 3). To perform this analysis, a symmetric rate matrix was assumed. To infer the intensity of directional transitions (forward and backward) within a matrix of the discrete traits mentioned above, we used a Markov jumps approach. To interpret the Bayes factors, a value of <3, as mentioned above, is not significant (hardly worth mentioning), BF= 3.1-20 represents positive support, BF= 20.1-150 represents strong support, while >150.1 represents an overwhelming support [75].

Finally, we visualized the spatiotemporal viral diffusion of each serotype by using Spatial Phylogenetic Reconstruction of Evolutionary Dynamics using Data-Driven Documents (D3) SPREAD3 software [76] considering the whole transmission and also the most significant connections between localities following the Bayesian stochastic search variable selection (BSSVS) method, with each country used as a discrete variable with a cutoff BF > 3 [75]. In addition, we classified the viral spread of each serotype in two categories: local, if the transmission occurs through neighboring countries, and long distance, if the dispersion jumps beyond adjacent neighboring countries.

## RESULTS

The number of available sequences per country varied from 1 to 18 whole genome sequences (WGS). Our results showed that India, China, Uganda, Argentina, and Zimbabwe were the countries with the highest number of available genomes (Supplementary Table S3). Likewise, several countries have been historically affected by more than one serotype, particularly in Africa and Asia, where we observed that Uganda presented the highest virus diversity as it has been subjected to the spread of serotypes O, and all SATs (Supplementary Table S3, Supplementary Fig. S1).

### Spatiotemporal dynamics of FMDV

Phylogeographic analyses highlighted great asymmetries in the tempo and mode of each serotype evolution (Fig. 1). SAT1 appeared to be the basal clade of the entire lineage, originating SAT2, SAT3 and serotype A, which later diversified into serotype Asia1, and O. Our analysis suggested O as the most recent, prolific and widespread lineage, with the highest number of sequences available worldwide. Serotype A and Asia1 appeared second and third in the number of available sequences, followed by all SAT serotypes. Maximum clade credibility phylogeny showed the monophyly of serotypes O, A, and Asia1 (each serotype shared a common ancestor), while SAT serotypes appeared to have multiple origins (Fig. 1).

**Fig. 1.**
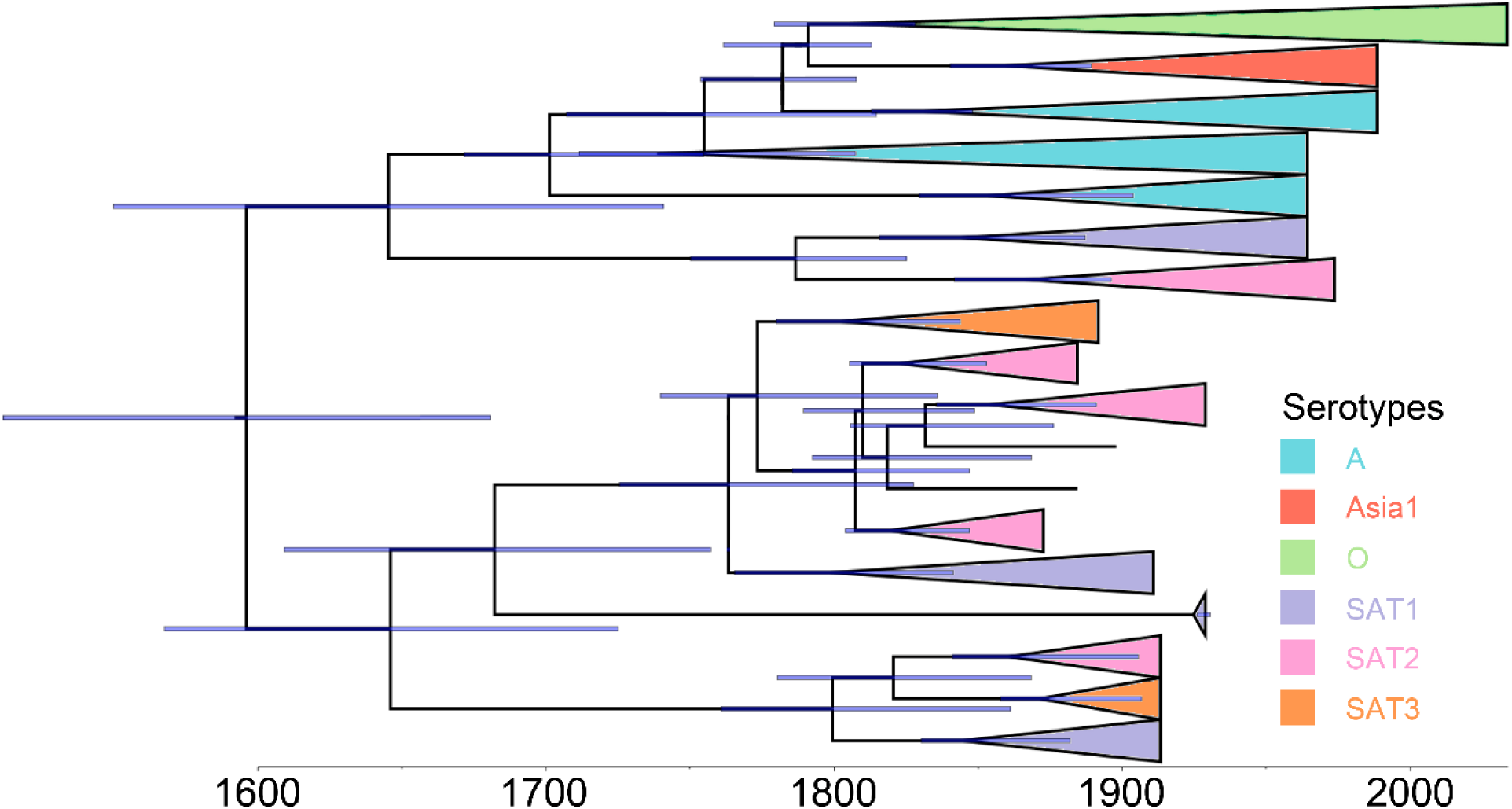
Condensed phylogenetic tree showing the overall evolutionary history of FMDV representing the relationships between all serotypes. The tree is based on a maximum clade credibility phylogeny inferred from 249 whole genome sequences. Branch bars represent posterior probabilities of branching events (*P* > 0.95).

### Spatiotemporal diffusion among serotypes

We investigated and compared the historical spreads of FMDV at serotype level, from the ones with local distribution (Asia1 and SATs) to the serotypes with widespread dispersal (O and A) (Sobrino et al., 2001; Fèvre et al., 2006; Di Nardo et al., 2011; Jamal and Belsham, 2013; Knight-Jones and Rushton, 2013; Brito et al., 2017).

### Serotype O

We analyzed 97 WGS from serotype O, which comprises 39% of the global FMDV tree (Fig. 1). This serotype also presents the widest distribution of all serotypes, with records from 42 countries (Fig. 2). Based on our phylogeographic analysis, the most likely center of origin for this serotype was Belgium (root state posterior probability [RSPP] = 0.27) from which it spread globally across long distances to several countries through Europe, Asia, Africa and South America. A remarkable aspect of this global spread is that most of it has occurred in less than 50 years (Fig. 3A) (see Supplementary Video S1 for detailed footage). These patterns have also appeared in our spatiotemporal diffusion map, which shows that the global distribution of this serotype is highly represented by long-range movements across countries and continents (Fig. 3A). Phylogenetic reconstruction identified clusters formed by different sub-lineages, where the most representative centers of dispersal events (geographic spread accompanied by diversification) for this serotype were Poland and the United Kingdom in Europe, China, Japan, and Indonesia in Asia, Egypt in Africa and Argentina in South America. Likewise, we observed that China, South Korea, and Turkey were also among the countries with the highest number of sequences (see Fig. 1A). In addition, BSSVS-BF results showed the most significant viral transmission routes for serotype O, where the most intense were represented from Turkey to Egypt, from Egypt to Indonesia and from Myanmar to Japan (BF>1038.7) (Fig. 3B).

**Fig. 2.**
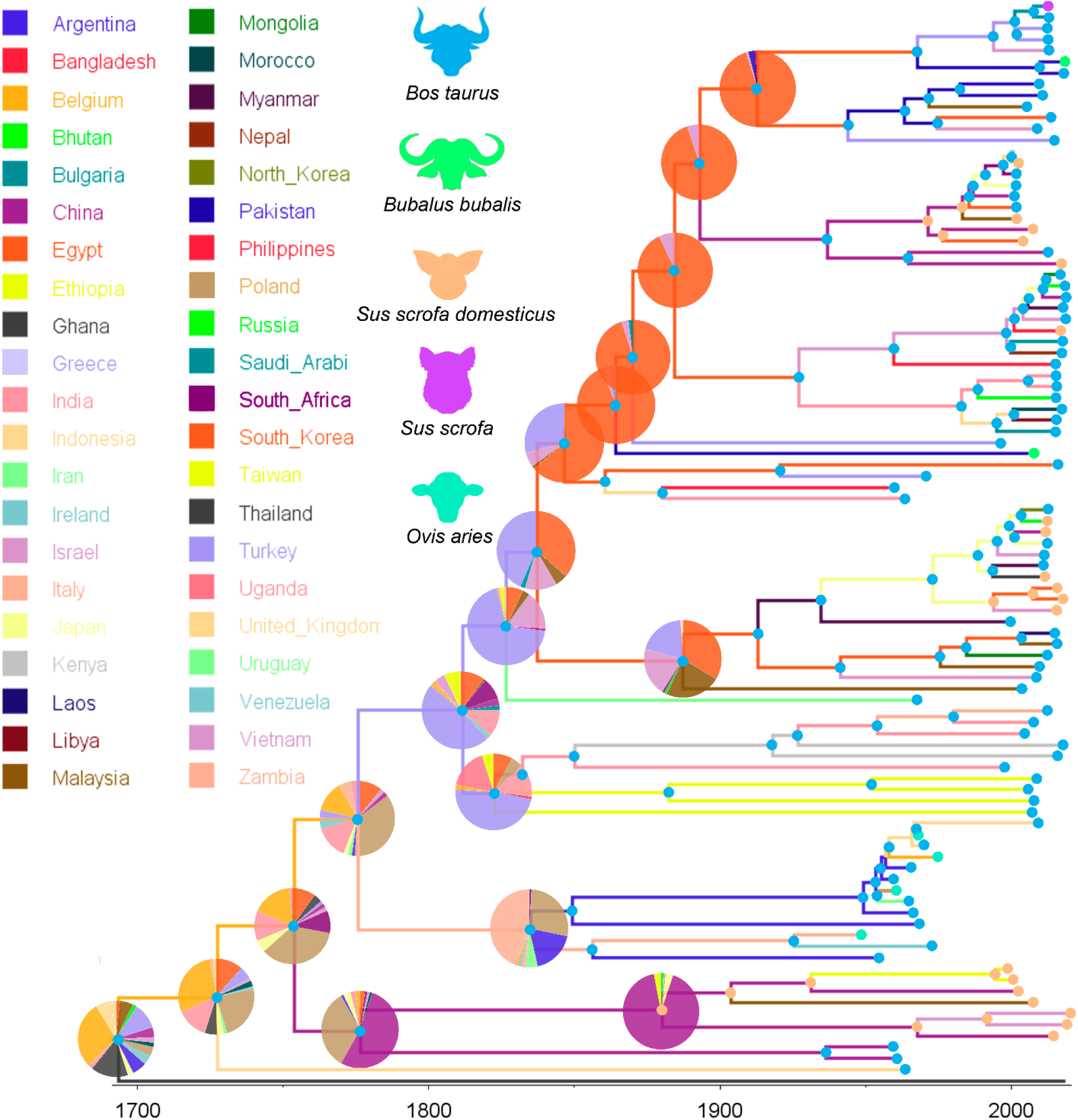
Dispersal history of FMDV lineages of Serotype O, as inferred by discrete phylogeographic analysis. Maximum clade credibility phylogeny colored according to the countries of origin. Branch bars represent posterior probabilities of branching events (P > 0.95). Colored dots at the end of the branches represent the host species (*Bos taurus=* cattle, *Sus scrofa domesticus*= swine, *Ovis aries=* sheep, *Bubalus bubalis=* water buffalo, and *Sus scrofa= boar*). The probabilities of ancestral states (inferred from the Bayesian discrete trait analysis) are shown in pie charts at each node, while circles on each branch and tips represent the most likely hosts.

**Fig. 3.**
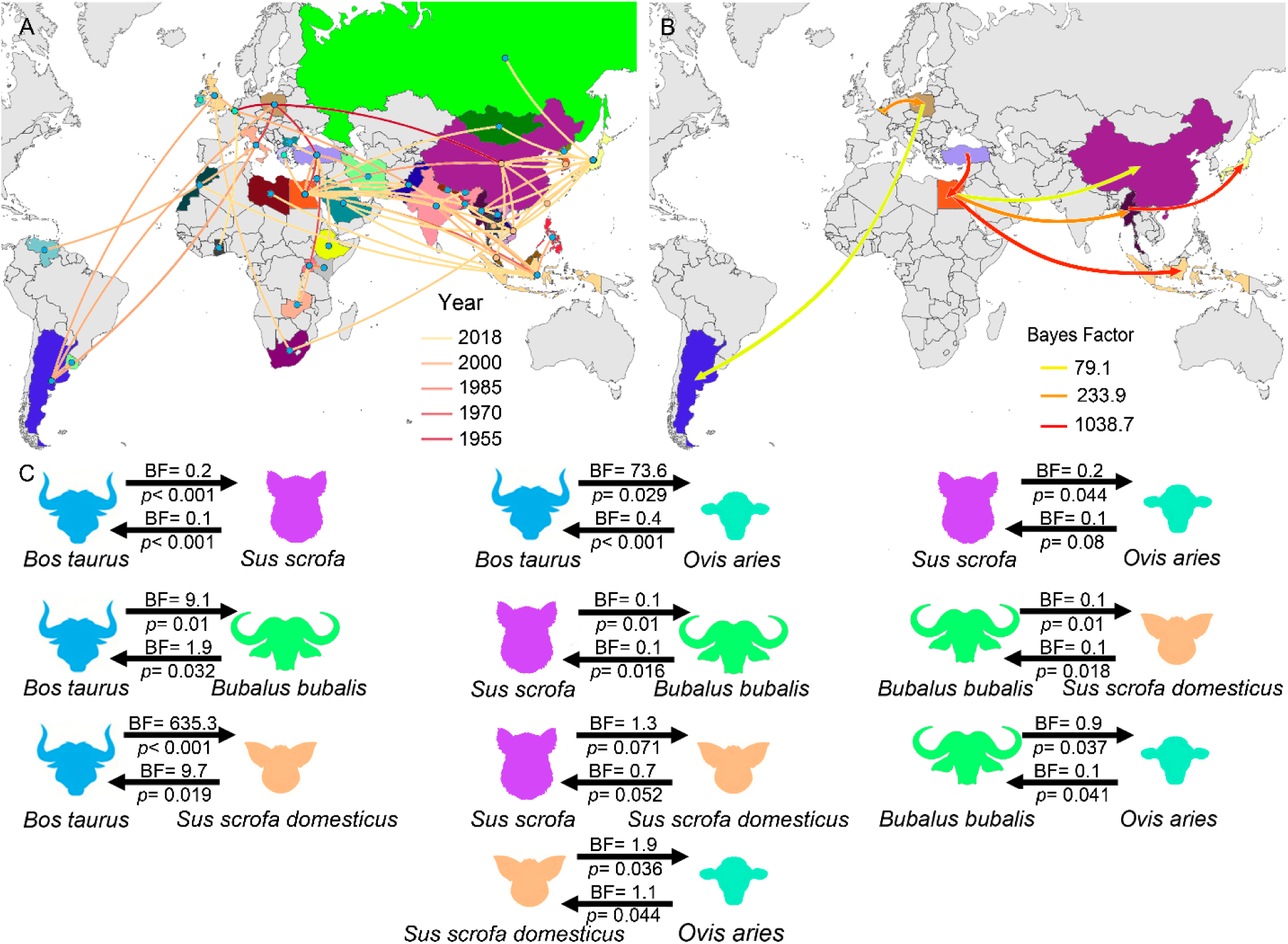
(A) Reconstructed spatiotemporal diffusion of FMDV serotype O spread, the color of the branches represents the age of the internal nodes, where darker red colors represent older spread events. (B) Representation of the most significant location transitions events for FMDV serotype O spread based on only the rates supported by a BF greater than 3 are indicated, where the color of the branches represent the relative strength by which the rates are supported. (C) Transmission rates between hosts (*Bos taurus=* cattle, *Sus scrofa domesticus*= swine, *Ovis aries=* sheep, *Bubalus bubalis=* water buffalo, and *Sus scrofa= boar*). based on BSSVS-BF values are represented on the top of the black arrows, while the root state posterior probability for the host-species transition are given on its bottom.

Serotype O also showed the highest host diversity among all FMDV serotypes, which is represented by cattle (*Bos taurus*), swine (*Sus scrofa domesticus*), boar (*Sus scrofa*), sheep (*Ovis aries*), and the water buffalo (*Bubalus bubalis*), where the most representative were *B. taurus* (71% of the sequences) and *S. scrofa domesticus* (20%). *B. taurus* was not only the most important host for this serotype but also the most likely initial host of the ancestral lineages (RSPP= 0.95), followed by *S. scrofa domesticus* (RSPP= 0.03) and *O. aries* (RSPP= 0.032) respectively (Fig. 2). Bayes factor analysis showed that the most significant transmission routes occurred from *B. taurus* to *S. scrofa domesticus* (BF= 635.3), followed by from *B. taurus* to *O. aries* (BF=73.6), and in a minor scale, from *S. scrofa domesticus* to *B. taurus* (BF= 9.7), as well as from *B. taurus* to *B. bubalis* (BF=9.1) (Fig. 3C).

The Bayesian skyline plot (BSP) was used to describe the observed changes in genetic diversity (population size) through time, showing a steady pattern in this serotype, with a sharp decrease in its effective population size occurred ∼2000, which returned to previous rates years later (Fig. S2).

### Serotype A

Phylogeographic relationships obtained for serotype A indicated India as its most likely center of origin (RSPP= 0.28, Fig. 4). Besides, several centers of diversification have been identified for this serotype, where the most important have been India and Malaysia in Asia, Netherlands and Germany in Europe, Chad in Africa and Brazil in South America (Fig. 4). Our phylogeographic analysis also highlighted the importance of Brazil as a center of origin for a wide variety of European and South American lineages (Fig. 4). As in serotype O, we observed that long-distance dispersal events were the most representative of the spatiotemporal dynamics of this serotype, evidencing a global distribution, with records from 33 countries (Fig. 5B), which represents 33% of the total FMDV sequences.

**Fig. 4.**
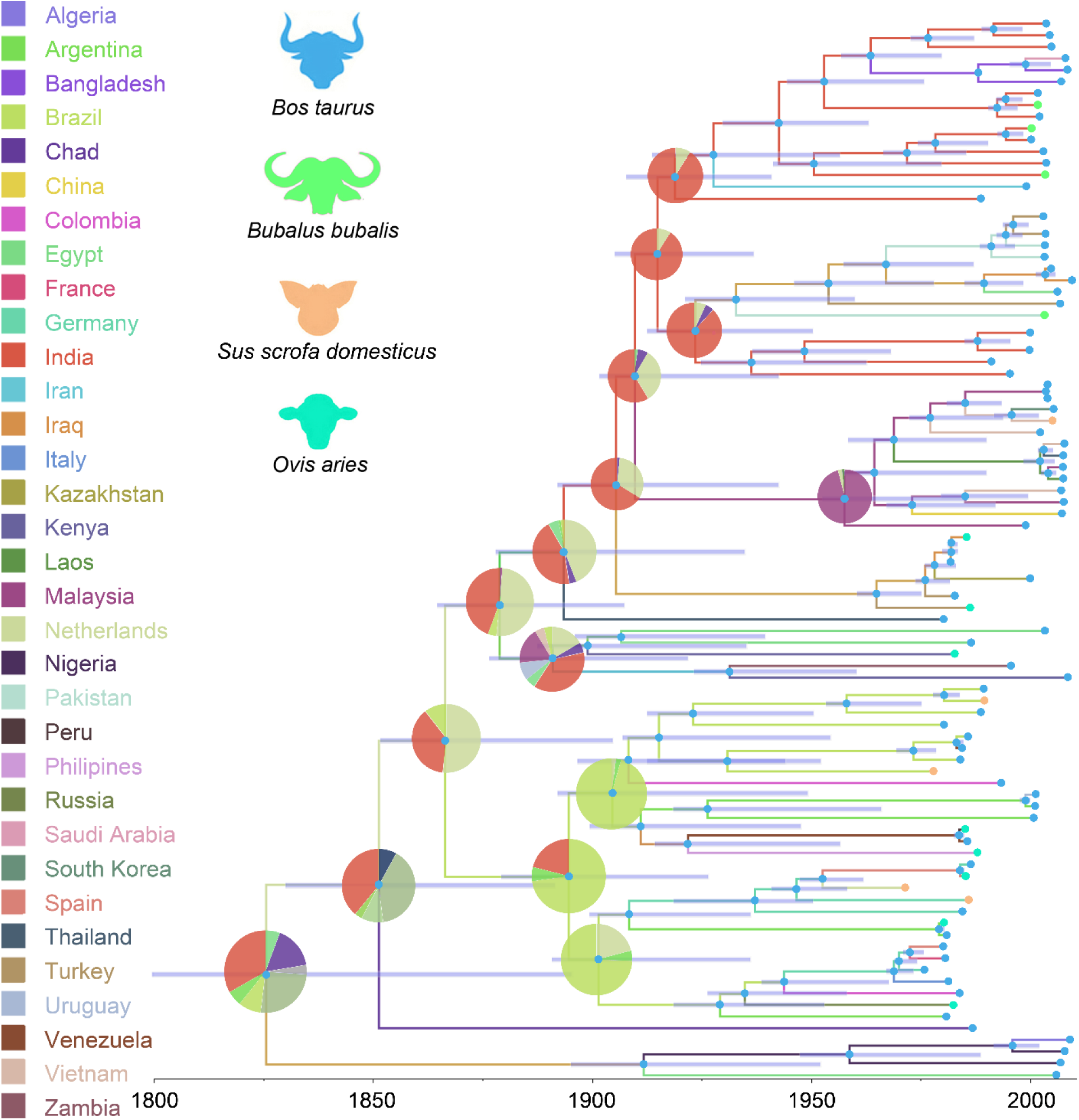
Dispersal history of FMDV lineages of Serotype A, as inferred by discrete phylogeographic analysis. Maximum clade credibility phylogeny colored according to the countries of origin. Branch bars represent posterior probabilities of branching events (P > 0.95). Colored dots at the end of the branches represent the host species (*Bos taurus=* cattle, *Sus scrofa domesticus*= swine, *Ovis aries=* sheep, and *Bubalus bubalis=* water buffalo. The probabilities of ancestral states (inferred from the Bayesian discrete trait analysis) are shown in pie charts at each node, while circles on each branch and tips represent the most likely hosts.

**Fig. 5.**
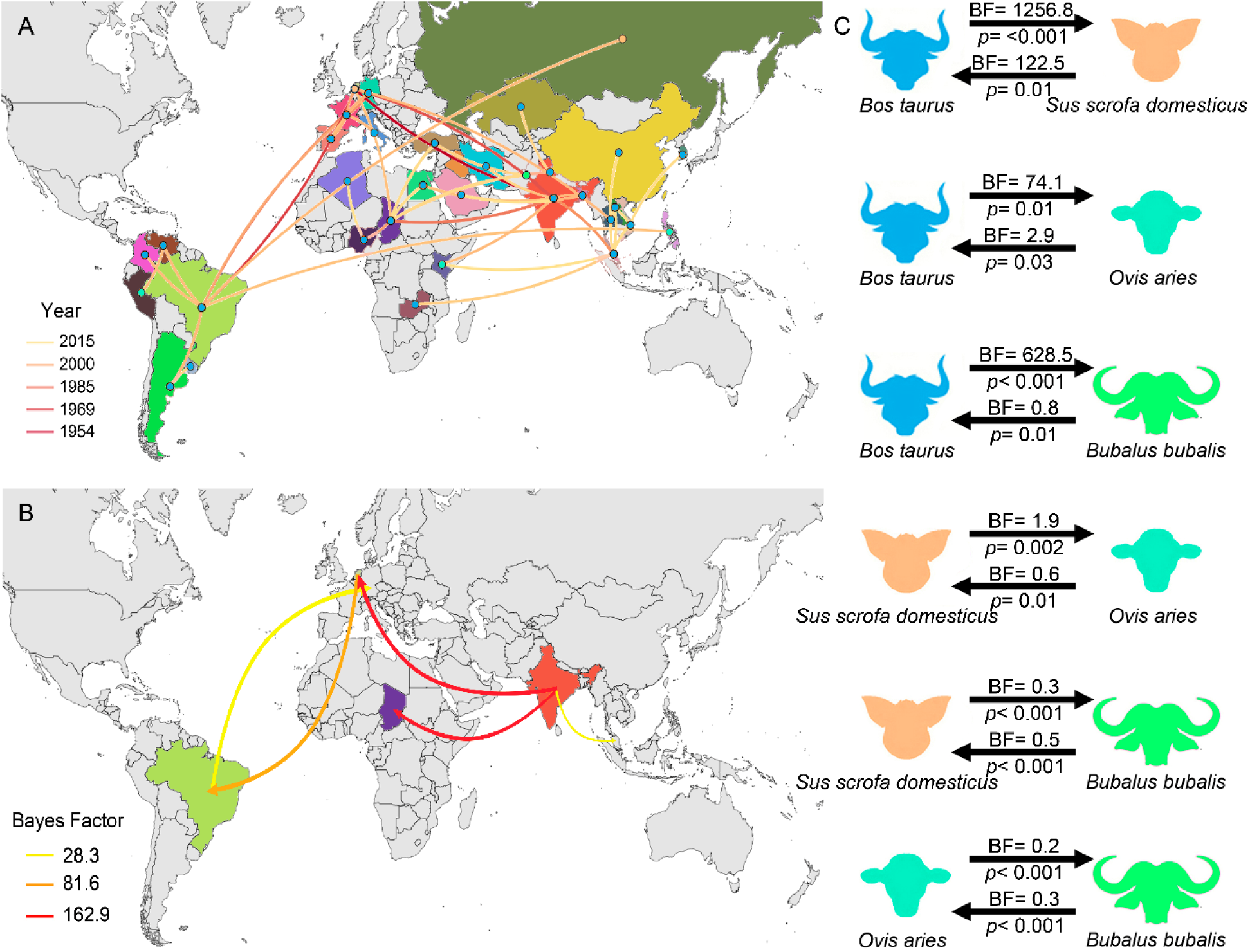
(A) Reconstructed spatiotemporal diffusion of FMDV serotype A spread, the color of the branches represents the age of the internal nodes, where darker red colors represent older spread events. (B) Representation of the most significant location transitions events for FMDV serotype A spread based on only the rates supported by a BF greater than 3 are indicated, where the color of the branches represent the relative strength by which the rates are supported. (C) Transmission rates between hosts (*Bos taurus=* cattle, *Sus scrofa domesticus*= swine, *Ovis aries=* sheep, and *Bubalus bubalis=* water buffalo) based on BSSVS-BF values are represented on the top of the black arrows, while the root state posterior probability for the host-species transition are given on its bottom.

BSSVS-BF analysis evidenced that the most significant transmission routes for this serotype come from India. Bayesian Factor analysis describing the most important viral transmission routes highlighted the importance of India in different directions, mainly to Netherlands (BF= 173.2), to Chad (BF= 162.9), and to Malaysia (BF= 28.3). Likewise, some European, such as Germany appeared to be important for the spread of this serotype into South America (BF=81.6, Fig. 5B).

The host species that showed highest number of serotype A sequences comes from *B. taurus* (82%), followed by *O. aries* (10%), *S. scrofa domesticus* (6%), and *B. bubalis* (2%) (Supplementary Table S1). Thus, the most important host behind the origin of the analyzed sequences of serotype A was *B. taurus* (RSPP= 0.98, Fig. 4). Furthermore, Bayes factor analysis indicated that the most significant transmission routes for the spread of this serotype occurred from *B. taurus* to all the other host. In order of significance, we can observe: to *S. scrofa domesticus* (BF= 1256.8), to *B. bubalis* (BF= 628.5), and to *O. aries* (BF= 74.1, Fig. 5C).

The BSP for this serotype showed a constant lineage diversity with a slight increase in 1980. The biggest variations between the years 2000 - 2017, showed a sharp decrease followed by a rapid increase in lineage diversity that was maintained until 2010 when these values mostly returned to the original values (Fig. S2).

### Serotype Asia1

This serotypes represented 12% of the entire FMDV sequences. Similarly to serotype A, phylogeographic analyses indicated India as the most likely origin of this serotype (RSPP= 0.34), from which it diverged in all directions (Fig. 6, Supplementary Video S3). The phylogenetic relationships seen in this serotype show a clear disparity between the lineages found in countries from western Asia (i.e., Israel, Lebanon, Afghanistan, and Turkey) and eastern Asia (i.e., China, Mongolia, Malaysia, and Vietnam) (Fig. 7A). Phylodynamic analysis shows that the most important centers of diversification for this serotype are India, China, Pakistan and Vietnam (Fig. 7A). BSSVS-BF analysis showed similar results as the observed in serotype A, where the most significant transmission routes are related to India. In order of intensity, the most important routes are the ones from India to China (BF= 182.8) and from India to Vietnam (BF= 61.5) (Fig. 7B).

**Fig. 6.**
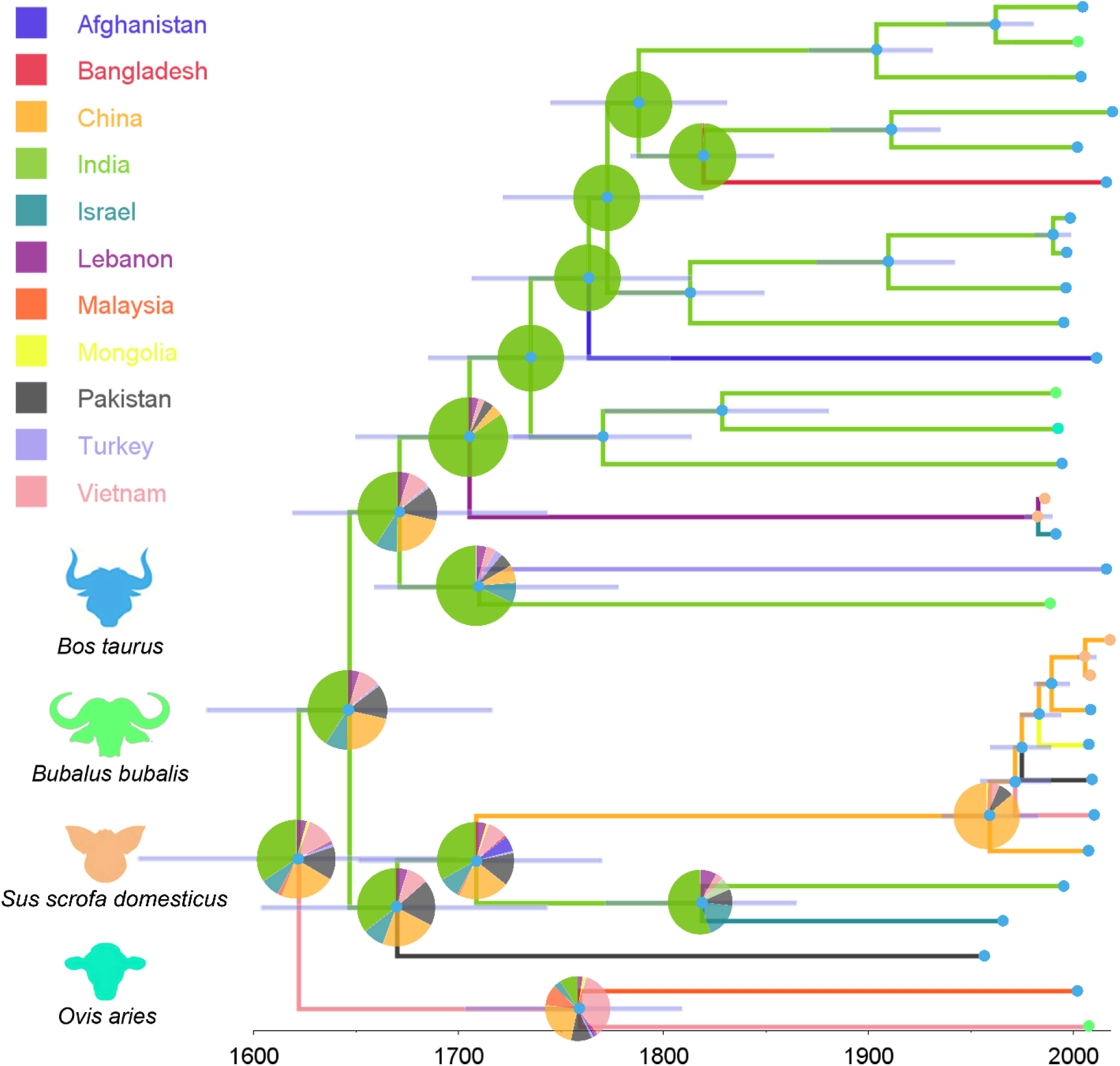
Dispersal history of FMDV lineages of Serotype Asia1, as inferred by discrete phylogeographic analysis. Maximum clade credibility phylogeny colored according to the countries of origin. Branch bars represent posterior probabilities of branching events (P > 0.95). Colored dots at the end of the branches represent the host species (*Bos taurus=* cattle, *Sus scrofa domesticus*= swine, *Ovis aries=* sheep, and *Bubalus bubalis=* water buffalo. The probabilities of ancestral states (inferred from the Bayesian discrete trait analysis) are shown in pie charts at each node, while circles on each branch and tips represent the most likely hosts.

**Fig. 7.**
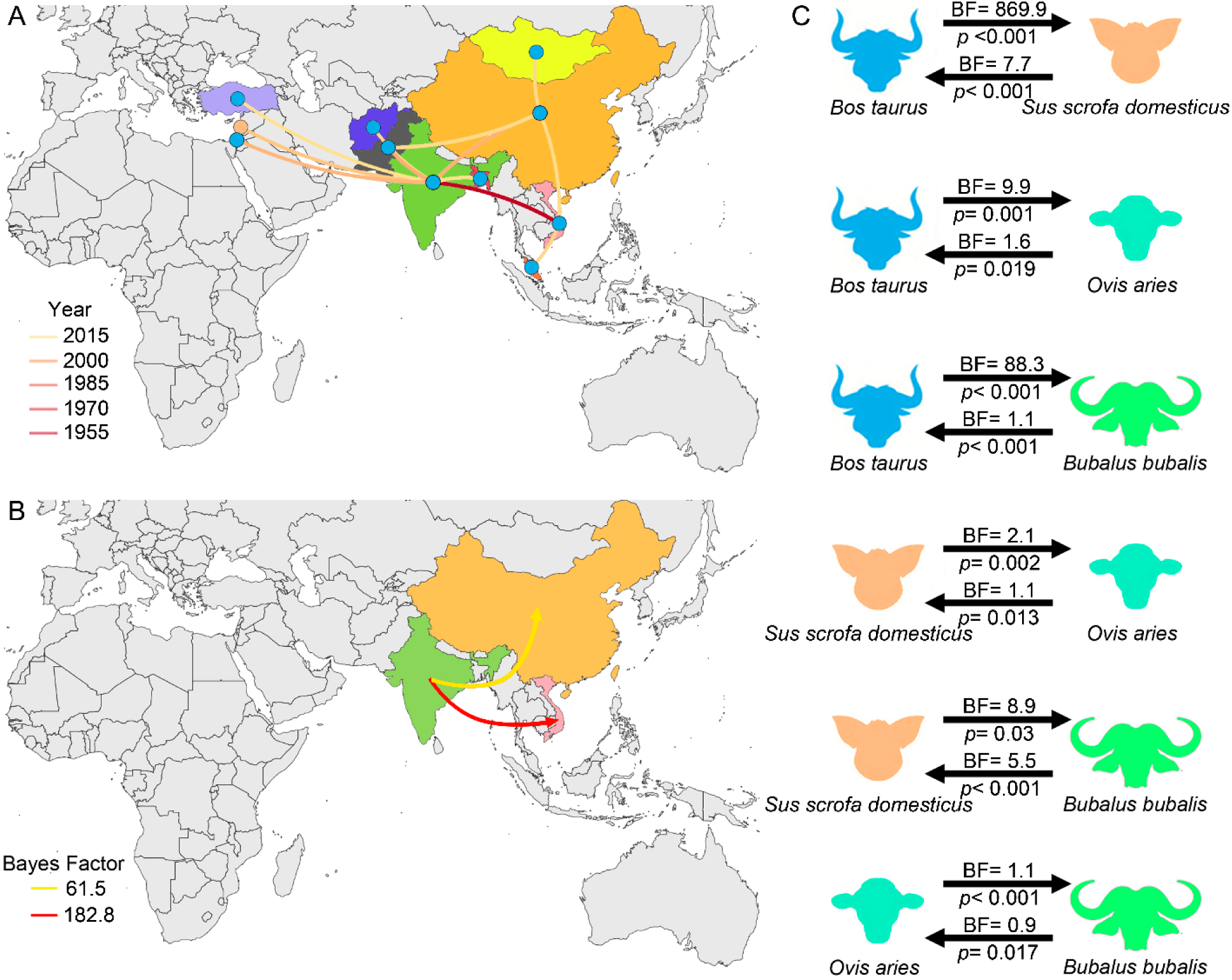
(A) Reconstructed spatiotemporal diffusion of FMDV serotype Asia1 spread, the color of the branches represents the age of the internal nodes, where darker red colors represent older spread events. (B) Representation of the most significant location transitions events for FMDV serotype Asia1 spread based on only the rates supported by a BF greater than 3 are indicated, where the color of the branches represent the relative strength by which the rates are supported. (C) Transmission rates between hosts (*Bos taurus=* cattle, *Sus scrofa domesticus*=swine, *Ovis aries=* sheep, and *Bubalus bubalis=* water buffalo) based on BSSVS-BF values are represented on the top of the black arrows, while the root state posterior probability for the host-species transition are given on its bottom.

In relation to the number of available sequences, the most representative hosts for this serotype were *B. taurus* (66% of the sequences), followed by *S. scrofa* (14%, exclusively in China), *Bubalus bubalis* (6%, observed in India, Vietnam, and China) and *Ovis aries* (∼3%, observed only in China). Phylogenetic analysis showed that the hosts responsible for the spread of this serotype were *B. taurus* (RSPP= 0.98), followed by *S. scrofa* (RSPP= 0.2). In addition, Bayes factor analysis indicated that the most strongly supported transmission routes for the spread of this serotype occurred from *B. taurus* to *B. bubalis* (BF= 1405.6), followed by from *B. taurus* to *S. scrofa* (BF= 401.3), and from *B. taurus* to *O. aries* (BF= 134.1) (Fig. 7C).

Phylodynamic patterns of serotype Asia1 spread through BSP approach showed a constant lineage diversity over time, with a very slight variation around the year 2000 (Fig. S2).

### Serotype SAT1

The phylogeographic patterns of SAT1 exposed Uganda as its most likely country of origin (RSPP= 0.45), from where it spread to Namibia, Nigeria, and Chad (Fig. 8). The phylogenetic relationships identified three main clusters, one of them represented by the ancestor of the lineages found in Uganda and Chad, other by the lineages located in the countries that are part of the southern area of spread (i.e., Botswana, Mozambique, Namibia, South Africa and Tanzania), and the last cluster represented by the lineages found in Nigeria, which also presented the sub-lineage with the most recent appearance. Contrary to the previous serotypes, SAT1 did not present a clear source (country) of dispersal events, although its spread was mostly concentrated on Eastern Africa (Fig. 9A). Our results showed that the highest proportion of its dispersal events occurred across long distance countries (representing 63% of the cases, See Supplementary Video S4 for detailed footage). The most significant dispersal routes were strongly related to Uganda, which in order of intensity, were seen to happen from Uganda to Nigeria (BF= 35.2), which seemed the most significant, followed by the transition from Kenya to Tanzania (BF= 26.5) (Fig. 9B).

**Fig. 8.**
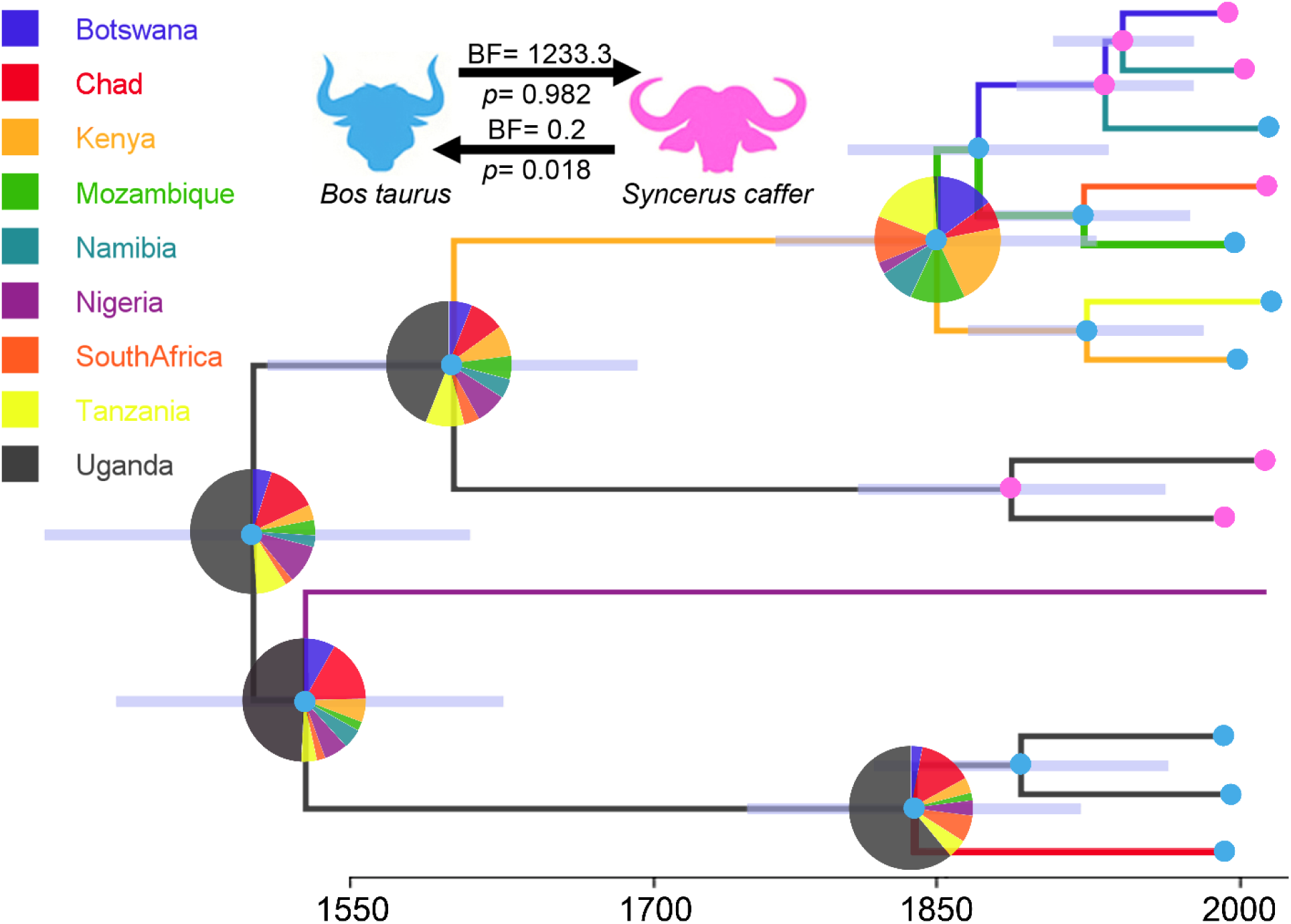
Dispersal history of FMDV lineages of Serotype SAT1, as inferred by discrete phylogeographic analysis. Maximum clade credibility phylogeny colored according to the countries of origin. Branch bars represent posterior probabilities of branching events (P > 0.95). Colored dots at the end of the branches represent the host species (*Bos taurus=* cattle, and *Syncerus caffer=* African buffalo. The probabilities of ancestral states (inferred from the Bayesian discrete trait analysis) are shown in pie charts at each node, while circles on each branch and tips represent the most likely hosts. Transmission rates between hosts (BSSVS-BF values) are represented on the top of the black arrows, while the root state posterior probability for the host-species transition are given on its bottom.

**Fig. 9.**
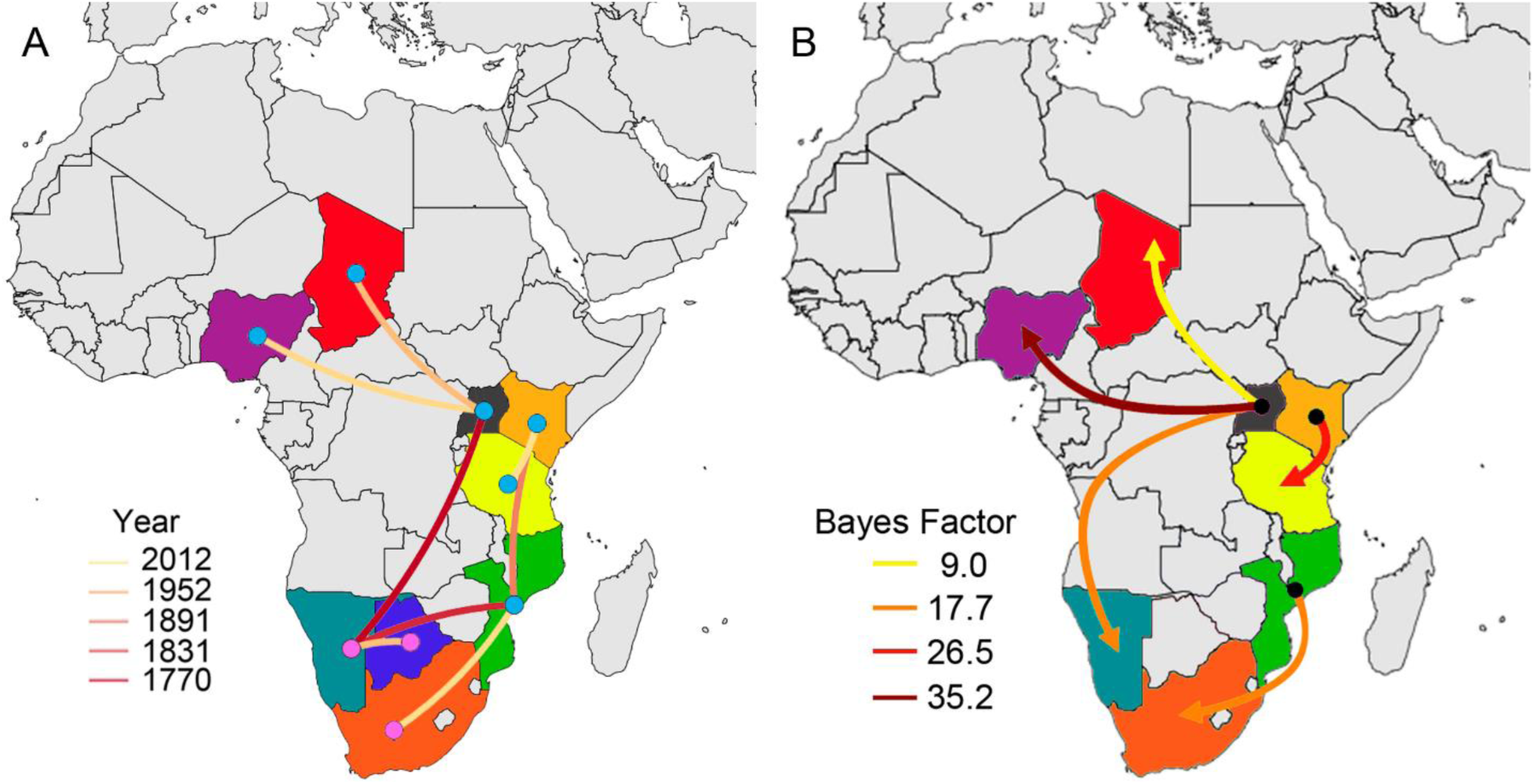
(A) Reconstructed spatiotemporal diffusion of FMDV serotype SAT1 spread, the color of the branches represents the age of the internal nodes, where darker red colors represent older spread events. (B) Representation of the most significant location transitions events for FMDV serotype SAT1 spread based on only the rates supported by a BF greater than 3 are indicated, where the color of the branches represent the relative strength by which the rates are supported.

The host species associated with the dispersal of this serotype were *Bos taurus* and *Syncerus caffer* (Fig. 8). *B. taurus*, with the majority of the number of sequences (58.3%), is mostly distributed in the north and central Africa, while *S. caffer* (41.7%, appeared as the most common host species in the southern countries (i.e., Namibia, Botswana, and South Africa). Phylogenetic analysis suggested *B. taurus* as the most important host species for the origin of this serotype (RSPP= 0.98). Strongly supported transmission routes were inferred from *B. taurus to S. caffer* (BF= 1233.3). However, the reverse transmission was not significant (BF<3) (Fig. 8).

Intriguingly, SAT1 BSP showed the most variable lineage diversity between all SATs, with an early increased in 1860 that was maintained until 1970, where this diversity increased again, reaching the values observed today (Fig. S2).

### Serotype SAT2

Phylogeographic analyses for SAT2 indicated Uganda as the most likely origin of the serotype (RSPP= 0.51), from which it spread to Botswana and The Gaza Strip later on time (Fig. 10). Later, from Botswana, this serotype expanded its distribution to Ethiopia, Zimbabwe, and Zambia, continuing spreading to surrounding countries also on the Eastern region of Africa (see Supplementary Video S5 for detailed footage). Phylogenetic analysis identified two main sub-lineages, one found between Uganda and Gaza Strip and the second cluster formed by the lineages found in countries distributed in southeastern Africa (Fig. 10). Likewise, our results also evidenced that serotype SAT2 spread is mostly characterized by a higher proportion of long-distance movements (57%) over local dispersal events (Fig. 11A). Our phylodynamic model suggested that the strongest geographic transition routes occurred from Uganda to Gaza Strip (BF= 34.6), and from Botswana to Zambia (BF= 23.2) (Fig. 11B).

**Fig. 10.**
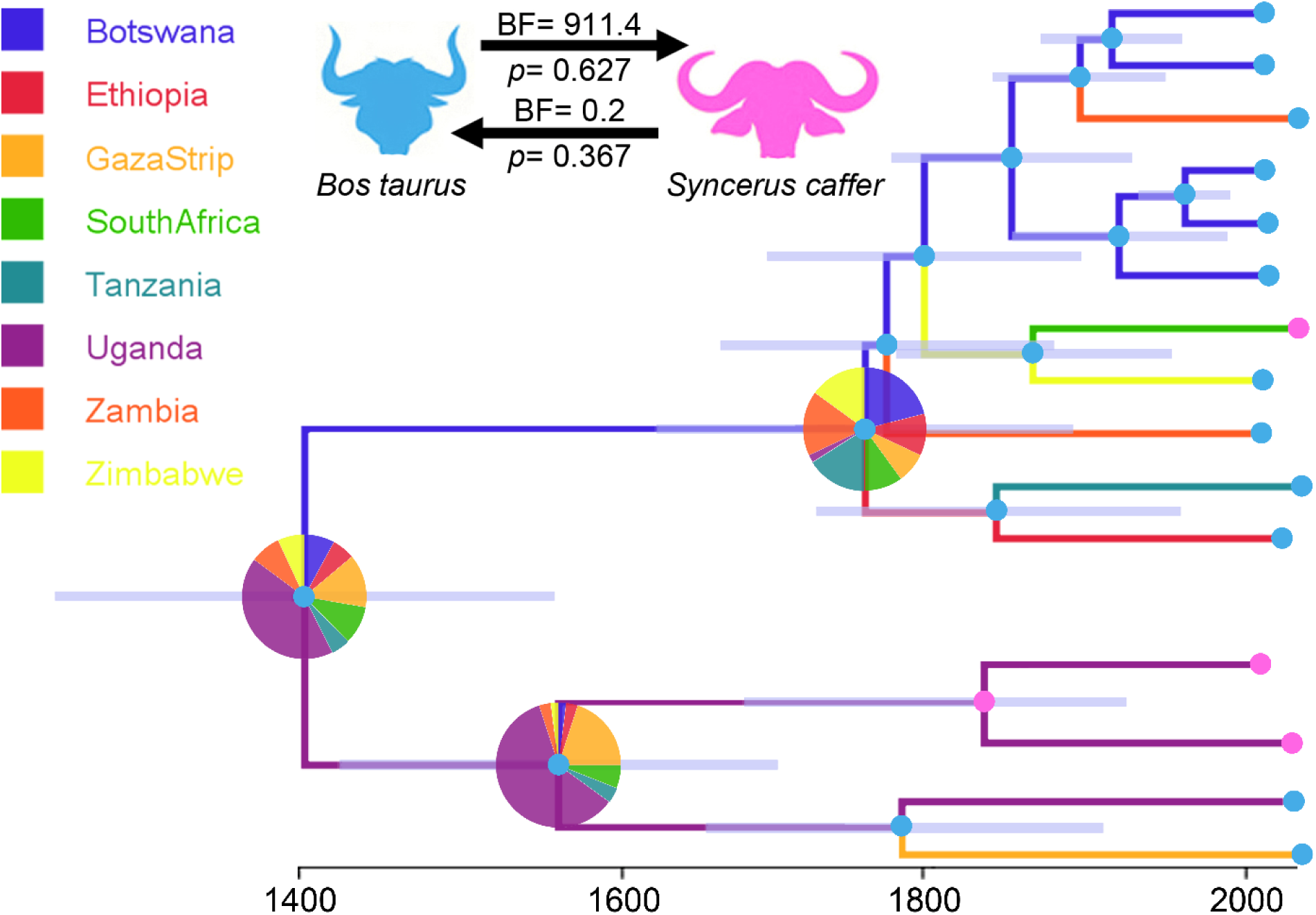
Dispersal history of FMDV lineages of Serotype SAT2, as inferred by discrete phylogeographic analysis. Maximum clade credibility phylogeny colored according to the countries of origin. Branch bars represent posterior probabilities of branching events (P > 0.95). Colored dots at the end of the branches represent the host species (*Bos taurus=* cattle, and *Syncerus caffer=* African buffalo. The probabilities of ancestral states (inferred from the Bayesian discrete trait analysis) are shown in pie charts at each node, while circles on each branch and tips represent the most likely hosts. Transmission rates between hosts (BSSVS-BF values) are represented on the top of the black arrows, while the root state posterior probability for the host-species transition are given on its bottom.

**Fig. 11.**
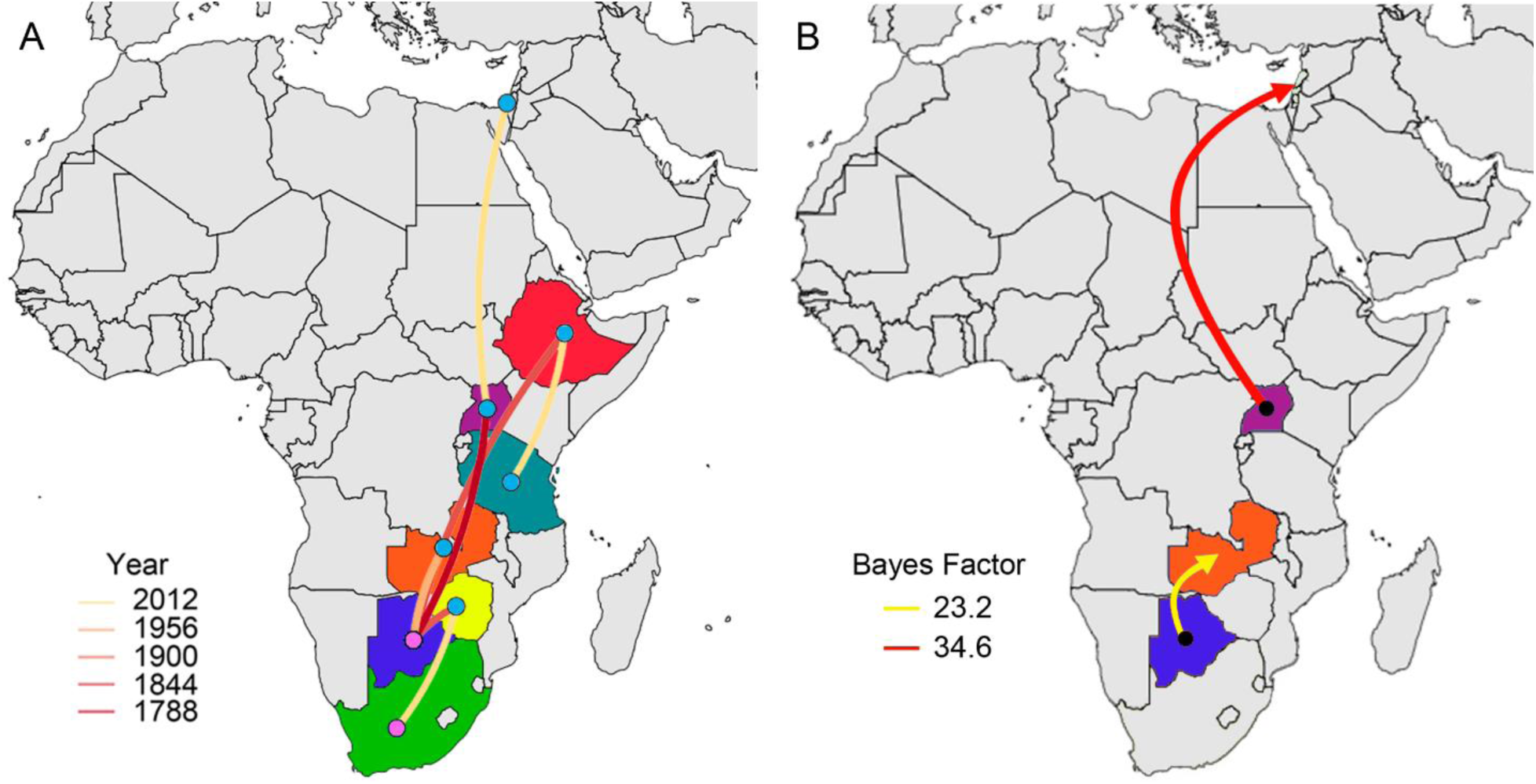
(A) Reconstructed spatiotemporal diffusion of FMDV serotype SAT2 spread, the color of the branches represents the age of the internal nodes, where darker red colors represent older spread events. (B) Representation of the most significant location transitions events for FMDV serotype SAT2 spread based on only the rates supported by a BF greater than 3 are indicated, where the color of the branches represent the relative strength by which the rates are supported.

*B. taurus and S. caffer* were also the main host associated with SAT2 sequences, but in this case, *S. caffer* was the most representative (53.9%), over *B. taurus* (46.1%). However, *B. taurus* appeared in most of the reported locations (except in South Africa), while *S. caffer* was only described in Uganda, Botswana and South Africa. As observed in SAT1, phylogenetic analysis showed a higher influence of *B. taurus* as the host of the ancestral lineages of this serotype (RSPP= 0.63) (Fig. 10). Transmission dynamics between host species suggested that transmission from *B. taurus to S. caffer* was the most important (BF= 911.4), while transmission from *S. caffer* to *B. taurus* was not significant (BF<3) (Fig. 7D). Finally, BSP showed no variation in the lineage diversity found over time (Fig. S2).

### Serotype SAT3

Similar to the previous SAT serotypes, SAT3 also had its ancestral origin in Uganda (RSPP= 0.49), from where it traveled to Zimbabwe and then spread to its neighboring countries (Fig. 12, see Supplementary Video S6). Phylogenetic analyses identified two main sub-lineages, one of them found in Uganda and the second (and most diverse), present in the countries that are part of southern Africa (i.e., Botswana, South Africa, Zambia, and Zimbabwe). Phylogeographic reconstruction reflected the importance of Zimbabwe for the spread of this serotype, being this country the most common center of origin for the diffusion of the disease to Zambia, Botswana and most recently to South Africa. Contrary to all the other serotypes (except for Asia1), spatiotemporal dynamics of serotype SAT3 showed that its spread has been dominated by local events (75%, Fig. 13A). Based on BSSVS-BF results, the most significant viral transmission routes for serotype SAT3 were represented by the dispersion from Zimbabwe to Botswana (BF= 15.9) and from Zimbabwe to South Africa (BF= 12.1) (Fig. 13B).

**Fig. 12.**
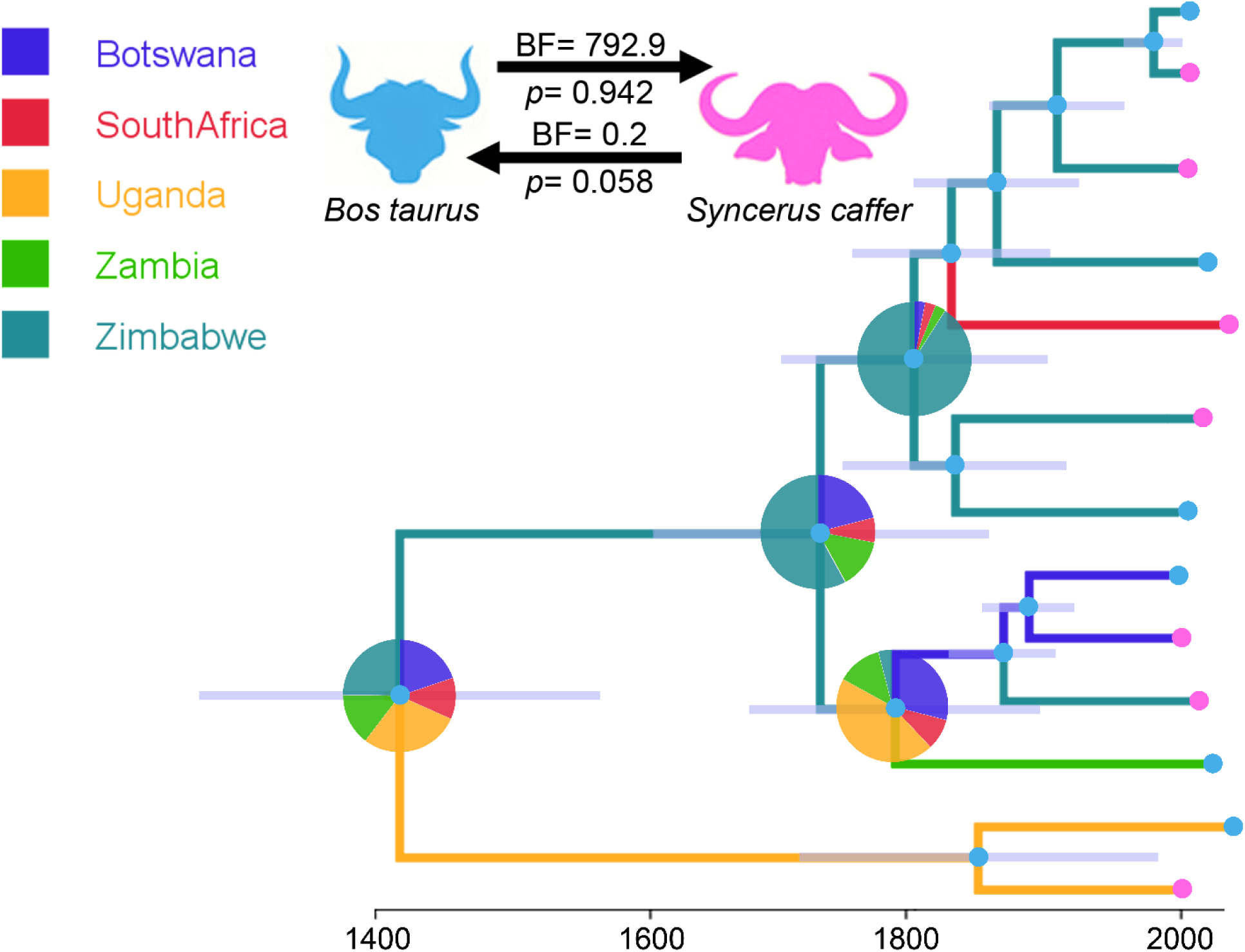
Dispersal history of FMDV lineages of Serotype SAT3, as inferred by discrete phylogeographic analysis. Maximum clade credibility phylogeny colored according to the countries of origin. Branch bars represent posterior probabilities of branching events (P > 0.95). Colored dots at the end of the branches represent the host species (*Bos taurus=* cattle, and *Syncerus caffer=* African buffalo. The probabilities of ancestral states (inferred from the Bayesian discrete trait analysis) are shown in pie charts at each node, while circles on each branch and tips represent the most likely hosts. Transmission rates between hosts (BSSVS-BF values) are represented on the top of the black arrows, while the root state posterior probability for the host-species transition are given on its bottom.

**Fig. 13.**
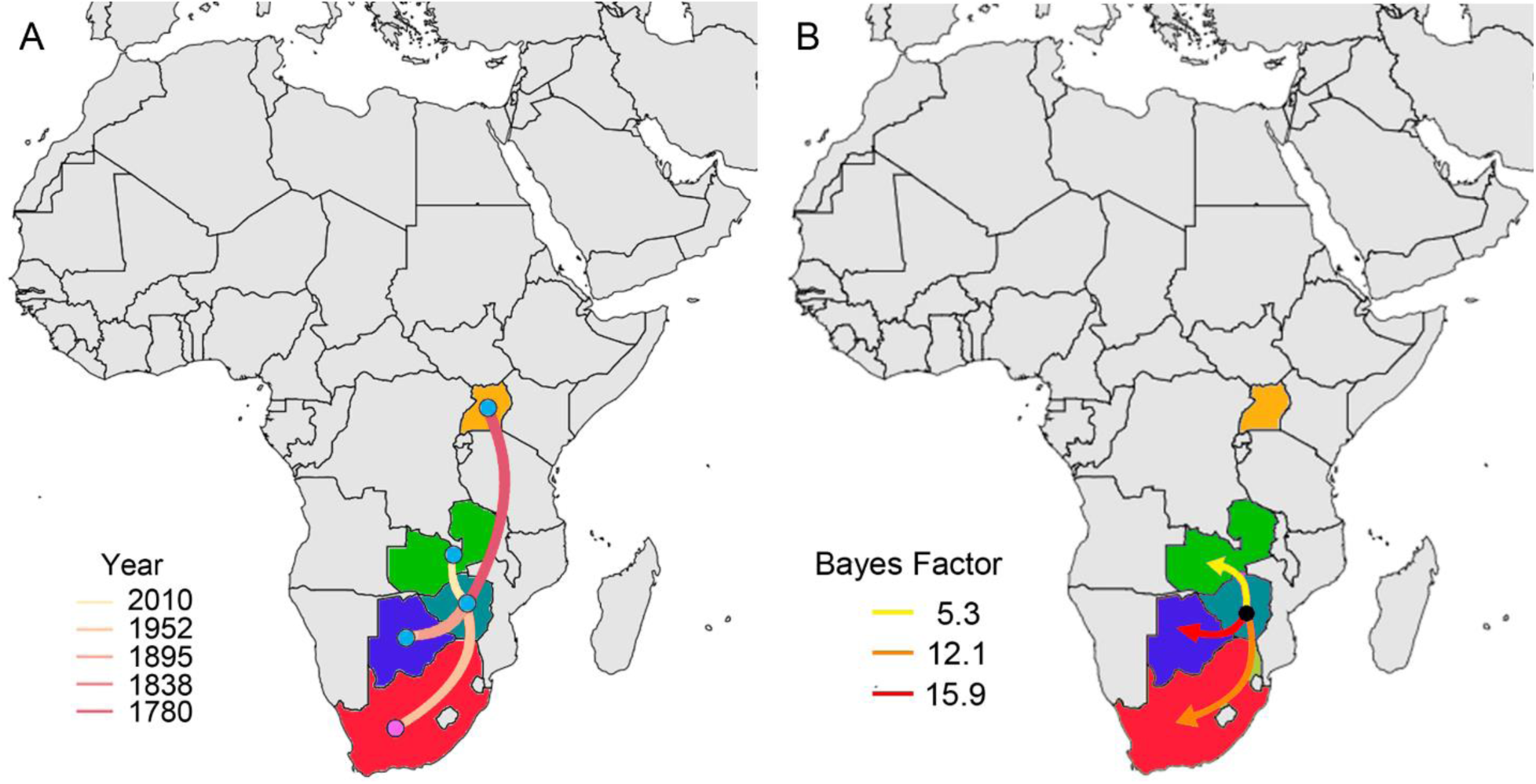
(A) Reconstructed spatiotemporal diffusion of FMDV serotype SAT3 spread, the color of the branches represents the age of the internal nodes, where darker red colors represent older spread events. (B) Representation of the most significant location transitions events for FMDV serotype SAT3 spread based on only the rates supported by a BF greater than 3 are indicated, where the color of the branches represent the relative strength by which the rates are supported.

*B. taurus and S. caffer* were also the main hosts reported for SAT3, appearing both in a similar proportion (*B. taurus*= 62.5%, and *S. caffer*= 37.5%). Similarly to all the serotypes above mentioned, cattle was the likely ancestral host species for this serotype (RSPP= 0.94). Furthermore, we found that most significant transmission routes for its spread occurred from *B. taurus to S. caffer* (BF= 792.9), while the transmission in the opposite direction was not significant (Fig. 12).

Like in the case of SAT2, the BSP obtained for this serotype showed no variation in the lineage diversity over time (Fig. S2).

## DISCUSSION

This study revealed new insights about the evolutionary dynamics of the FMDV’s global transmission dynamics at serotype level. The most likely country of origin for each serotype was identified, along with its historical spread characteristics, and divergence patterns across its historical dispersal. Finally, we assessed the impact of each host interaction in the spread of FMDV, providing a comprehensive characterization of transmission dynamics between host species.

### Phylogeographic patterns of FMDV spread

Global patterns of FMDV spread were considerably asymmetric in its spatiotemporal arrangement, showing important variation among all serotypes, as previously observed by Yoon et al., (2011) and Brito et al., (2015). On the other hand, our results yielded discrepancies regarding the phylogenetic relationships of FMDV serotypes due to the disagreements observed in the cladistic characterization of FMDV serotypes (monophyletic or polyphyletic origin) [52]. Lewis-Rogers et al. (2008) and Yoon et al. (2011) suggested that O, A, Asia1, C, and SAT3 were monophyletic, while SAT1 and SAT2 serotypes were polyphyletic. However, our results indicated the presence of only three monophyletic serotypes (O, A, and Asia1), whilst all SAT serotypes appeared to have multiple ancestral origins which can be related to multiple points of independent introduction of the virus.

#### Serotypes with global distribution (O and A)

Serotype O has shown a remarkable widespread distribution across the globe. In half of a century, this serotype reached almost all continents, causing dramatic economic losses [58, 77, 78]. Root state posterior probability analysis inferred Belgium as the most likely center of origin for this serotype, which, as a result of being responsible for the majority of outbreaks worldwide [79], we can observe multiple centers of diversification in most of the continents. Our phylogeographic analysis showed that this spread has been characterized by lineage dispersal events between distant regions (i.e. to regions not sharing international dry borders with the origin country), instead of dispersal events between neighboring countries, which may be one of the keys for its successful global spread. Bayesian skyline plot showed a severe decline in the genetic diversity around early 21^th^ century, this interesting pattern also observed by Yoon et al., (2011). This decline and recover in the effective population size could be directly related to the increase in the FMDV outbreaks that occurred worldwide, which was followed by an intensive control and prevention strategies. The intense wave of outbreaks occurred during that period worldwide included countries such as, Argentina (Perez et al., 2004; Perez et al., 2004), the United Kingdom [82, 83], Brazil [84], India [85], and Taiwan [86, 87]. As we observed in our phylogeographic visualization, there is strong evidence that most of these outbreaks were strongly interconnected [25, 88–90], evidencing local and long-distance spread of serotype.

One of the reasons for the success of the evolutionary diversification of this serotype may also be related to the diversity of hosts that it affects, which is the highest among all serotypes. Globally, *B. taurus* represented the most important host species for the spread of serotype O, while *S. scrofa* was mostly related to the spread of this serotype in southeastern Asia. Thus, phylodynamic analysis suggested that viral transition rate between these two livestock was the strongest reported between all the reported hosts.

Following the pandemic patterns showed by serotype O, the next large-scale potential of diffusion was exhibited by serotype A. Phylogeographic analysis suggested India as the most likely center of origin of the current circulating serotype A strains. Supporting previous studies [26, 60], we observed that India was also a key source of dispersal events for this serotype since most of the current strains are strongly related to India. Whole genome sequences of this serotype have been recorded in three continents, Asia, Africa, and South America, where it was reported as the causing agent of one of the biggest FMDV outbreaks, which occurred in Argentina in 2011, affecting a total of 2,126 herds [81]. It is important to note, that there is evidence of a posterior spread of these serotypes (O and A) to other countries, mainly in South America, since both of them are currently found in nearly every country of the continent [26, 79]. However, due to the lack of whole-genome data, we were unable to further assess this spread.

As expected, the main host affected by serotype A was *B. taurus*. This species had an important role in its viral spread [56], especially in this globalized era, where the continuous increase in livestock trading markets facilitates the spread of transboundary animal diseases [91]. Likewise, our phylodynamic analysis showed that the most intense host species transmission route occurred from *B. taurus* to *S. caffer*, and apparently, the reverse transmission is an infrequent event.

#### Asia1 and SAT serotypes

Whereas our results showed that serotypes O and A have spread worldwide, serotypes Asia1 and SATs remained non-pandemic and confined in their endemic regions [79, 92]. Since there is a lack of detailed sequences data available, especially for African countries, it is important to note that these results may vary with a better representation of the currently circulating virus, although they support what has been previously described [59, 79, 93, 94].

Undoubtedly, India has been historically considered as one of the most important countries for the spread and maintenance of FMDV, especially for serotypes A, and Asia1 [26, 55, 79, 94] Indeed, our phylogeographic analyses showed India as the most likely origin country for Asia1 serotype [26, 94]. The spread of this serotype was mainly restricted to Asia [53, 55, 93], and characterized by local movements across the neighboring countries surrounding India, China and Malaysia, where it is well known that free and unrestricted animal movements across country borders may play a key role in the spread of FMDV [55, 95]. We also observed India as a key center of dispersal for this serotype, which coincides with previously reported results [55]. The arrival of Asia1 into Turkey in 2013 represents one of the most recent and longer dispersal events reported for this serotype, which was directly related to an Indian sub-lineage of the virus [96]. Likewise, there have been sporadic incursions into other countries such as Greece in 1984 and 2000 [93], Malaysia in 1999 [31] or Turkey in 2017 [55], whose outbreaks seemed to be caused also by independent sub-lineages from the rest of the outbreaks observed in these regions.

Despite a previous study described multiple potential origins for SAT serotypes, (i.e., SAT1 in Zimbabwe and SAT2 in Kenya [48]), our root state posterior probability results suggested Uganda as the most likely origin for all of them. Likewise, our phylogeographic analysis also highlighted the importance of Uganda as a primary source of dispersal events to different countries, where the most strongly significant routes were found from Uganda to Nigeria (SAT1), from Uganda to Gaza strip (SAT2) and from Zimbabwe to Botswana (SAT3).

SAT serotypes (SAT1, SAT2, and SAT3) are characterized by a higher proportion of local spread, limited across their endemic areas. This spread occurred mainly in southeastern Africa, where nomadic pastoralism across international borders and animal trade in the sub-Saharan region is one of the most practiced forms of livestock movements [48, 56, 93, 94]. These results complement the observations made by Bouslikhane (2015), who highlighted how nomadism and transhumance play a key role in disease transmission, especially in African countries.

Previous studies have highlighted the importance of African buffalo (*Syncerus caffer*), hypothesizing that current FMDV genotypes may emerge in domesticated host species from viral reservoirs maintained by this species [49, 53, 59, 94, 98–104] However, the uncertainty over the involvement of African buffalo arose the need for deeper research to confirm its influence in livestock outbreaks [94]. Our results coincide with the evidence mentioned in a recent study by Omondi et al (2019), where cattle appeared as the most important host species for the spread of FMDV, while buffalo played a secondary role. This pattern was observed not only in SATs but in all serotypes studied.

In general, we observed considerable differences in the spatiotemporal dynamics exhibited by the different serotypes. Where the serotypes with global distribution (O and A) presented the most asymmetrical pattern in the annual genetic diversity in comparison with (SAT and Asia1 serotypes). Cattle was observed to play a key role in the historical spread of all serotypes of FMDV. Likewise, our phylodynamic analysis inferred that the transmission route from cattle to buffalo was the most highly supported, pattern that was also observed for all serotypes, independently of its spread potential.

Serotypes such as Asia1 and SATs presented local spread rates, mainly associated with cattle and sheep (with special importance of buffalo in the case of SATs serotypes) supporting previously described results (Brito et al., 2015; Omondi et al., 2019), while serotypes O and A showed long-distance spread, covering higher extensions of territory between each outbreak, which also confirms previously described information [59]. These serotypes presented the highest variety of susceptible hosts, although we speculate that the main reason for their successful long-distance spread relies mostly on the international movement of cattle and swine due to the intensive commerce between countries.

Finally, important limitations related to the use of whole genome relay in the lack of good global data, especially in African countries which remains endemically affected by five different serotypes, therefore some countries with known FMDV circulation are not part of this study. However, to reduce the bias generated by the strong unbalance of the available data in both dimensions (i.e., number of samples per country and uneven number of samples per host species), we removed all the sequences that where duplicated (i.e., represented the same outbreak multiple times), which, in the case of big outbreaks such as United Kingdom 2001/2007, Argentina 2001, and Japan 2010, accounted for hundreds of sequences representing each event. This limitation is common among phylogenic studies with no yet best alternative, this is true mainly because sample that are available hosted in public databases or from diagnostic laboratories [105]. Although whole genome sequences are increasingly proving to be a more accurate tool for phylogenetic analyses [106, 107], its high cost in comparison to studies considering partial genome results in lower availability of WGS, which became the major limitation for the construction of our dataset, resulting several countries with known reports of FMDV but lacking genetic data. Finally, it is important to highlight that, despite the nucleotide sequences encoding the capside protein VP1, VP2, and VP3 are sufficient to identify FMDV at serotype level, we preferred using WGS because of its higher accuracy in the determination of the genetic relationship between the reported cases [108].

### Final remarks

Studies considering whole genome sequences should be preferred over partial sequence research to ensure the importance of considering virus spread in its overall context [53, 106, 107]. Besides, the growing awareness of the importance of using whole genome sequences to assess the evolution of infectious diseases, and more specifically for RNA viruses as FMDV plays a key role on the future ability to analyze the ever-increasing volume of data accurately, getting closer to a real-time assessing of disease outbreaks [106]. However, the use of whole genome sequences represented a limitation in our study since the lack of FMDV sequences in a given country does not mean that the virus has not been circulating in that country but maybe associated with technical or economic constraints, therefore interpretation requires caution due to the possible introduction of sampling bias. The popularization of whole genome sequencing will help not only to increase the available information about the virus, but also have a direct impact on promoting new and more specific measures for disease control [24, 59, 109]. The result of such improvement in disease surveillance would not only be beneficial for the targeted region, but also for all the areas that are directly connected (i.e., through geographical limits) and indirectly (i.e., through commercial networks), including countries currently considered as free zones [110].

## CONCLUSION

In summary, we have seen how FMDV evolved and diversified in five species among 64 countries, by using a comprehensive phylodynamic approach, we characterized and compared its global phylogeographic distribution at serotype scale. The phylogeographic approach used here relies on the principle that evolutionary processes are better understood when a broader spatiotemporal vision is available. Our results shed light on FMDV’s macroevolutionary patterns and spread, allowing to unravel the ancestral country of origin for each serotype as well as the most important historical routes of viral dispersal, the role that the main host species played in its spatial diffusion and how likely the disease is transmitted between them. The use of whole genome sequences allowed us to clarify past discrepancies related to the polyphyletic nature of some serotypes (i.e., SATs), previously described as monophyletic.

Based on our findings, we corroborate with recent advancements that have been undertaken to control global distribution of major arbovirus (i.e., Dengue, yellow fever and Zika) [111–113], with the need to also implement real-time genome-scale sequencing to food-animal epidemics, in which metagenomics and phylogeography approaches inform epidemic responses and improve control intervention strategies.

## ACKNOWLEDGEMENTS

We acknowledge the Department of Population Health and Pathobiology-North Carolina State University provided startup funds for G. Machado and M. Jara. SD is supported by the Fonds National de la Recherche Scientifique (FNRS, Belgium). Guy Baele acknowledges support from the Interne Fondsen KU Leuven / Internal Funds KU Leuven under grant agreement C14/18/094.

## SUPPLEMENTARY MATERIAL

**TABLE S1.** Sample information for all Foot and Mouth Disease virus complete genome sequences used in this study.

**TABLE S2.** Root-to-tip regression analyses of phylogenetic temporal signal. Correlation and determination coefficient (R2) were estimated with TempEst (Rambaut et al. 2016). P-values were calculated using the approach of Murray et al. (2016) and were based on 1,000 random permutations of the sequence sampling dates (Navascuès et al. 2010).

**TABLE S3.** Number of sequences and serotypes per country.

**Fig. S1** Spatial distribution of Foot-and-mouth disease virus showing the number of serotypes per country.

**Fig. S2** Reconstructed Bayesian Coalescent Skyline plots (BSP) of FMDV serotypes. The median estimated of the effective population size through time are represented by the dark blue. The 95% highest posterior density confidence intervals are marked in blue.

**Video S1.** Reconstructed spatiotemporal diffusion of FMD serotype O spread, where diameters of the colored circles are proportional to the square root of the number of MCC branches, maintaining a particular location state at each time period. The color of the branches represents the age of the internal nodes, where darker red colors represent older spread events, this visualization match with the main time bar on top of the video.

**Video S2.** Reconstructed spatiotemporal diffusion of FMD serotype A spread, where diameters of the colored circles are proportional to the square root of the number of MCC branches, maintaining a particular location state at each time period. The color of the branches represents the age of the internal nodes, where darker red colors represent older spread events, this visualization match with the main time bar on top of the video.

**Video S3.** Reconstructed spatiotemporal diffusion of FMD serotype Asia1 spread, where diameters of the colored circles are proportional to the square root of the number of MCC branches, maintaining a particular location state at each time period. The color of the branches represents the age of the internal nodes, where darker red colors represent older spread events, this visualization match with the main time bar on top of the video.

**Video S4.** Reconstructed spatiotemporal diffusion of FMD serotype SAT1 spread, where diameters of the colored circles are proportional to the square root of the number of MCC branches, maintaining a particular location state at each time period. The color of the branches represents the age of the internal nodes, where darker red colors represent older spread events, this visualization match with the main time bar on top of the video.

**Video S5.** Reconstructed spatiotemporal diffusion of FMD serotype SAT2 spread, where diameters of the colored circles are proportional to the square root of the number of MCC branches, maintaining a particular location state at each time period. The color of the branches represents the age of the internal nodes, where darker red colors represent older spread events, this visualization match with the main time bar on top of the video.

**Video S6.** Reconstructed spatiotemporal diffusion of FMD serotype SAT3 spread, where diameters of the colored circles are proportional to the square root of the number of MCC branches, maintaining a particular location state at each time period. The color of the branches represents the age of the internal nodes, where darker red colors represent older spread events, this visualization match with the main time bar on top of the video.

